# Categorizing prediction modes within low-pLDDT regions of AlphaFold2 structures

**DOI:** 10.1101/2025.06.06.658382

**Authors:** Christopher J Williams, Vincent B Chen, David C Richardson, Jane S Richardson

## Abstract

AlphaFold2 protein structure predictions are widely available for structural biology uses. These predictions, especially for eukaryotic proteins, frequently contain extensive regions predicted below the pLDDT 70 level, the rule-of-thumb cutoff for high confidence. This work identifies major modes of behavior within low-pLDDT regions through a survey of human proteome predictions provided by the AlphaFold Protein Structure Database. The *near-predictive* mode resembles folded protein and can be a nearly accurate prediction. *Barbed wire* is extremely unproteinlike, being recognized by wide looping coils, an absence of packing contacts, and numerous signature validation outliers, and it likely represents a nonpredicted region. *Pseudostructure* presents an intermediate behavior with a misleading appearance of isolated and badly formed secondary structure-like elements. These prediction modes are compared with annotations of disorder from MobiDB, showing general correlation between *barbed wire/pseudostructure* and many measures of disorder, an association between *pseudostructure* and signal peptides, and an association between *near-predictive* and regions of conditional folding. To enable users to identify these regions within a prediction, a new Phenix tool is developed encompassing the results of this work, including prediction annotation, visual markup, and residue selection based on these prediction modes. This tool will help users develop expertise in interpreting difficult AlphaFold predictions and identify the near-predictive regions that can aid in molecular replacement when a prediction does not contain enough high-pLDDT regions.

## 1. Introduction

AlphaFold structure prediction (Jumper et al., 2021) has revolutionized structural biology experimental design (Thornton et al., 2021) and greatly eased the early stages of experimental structure solutions (Terwilliger et al., 2023) (Terwilliger et al., 2024). High-confidence AlphaFold predictions make excellent molecular replacement targets for x-ray crystallography and starting models for cryoEM. AlphaFold’s own pLDDT (predicted Local Distance Difference Test) provides a guide to whether a region is usable for structural biology, with pLDDT >= 70 serving as a rule of thumb cutoff for high confidence regions, and pLDDT >= 90 for very high confidence. However, especially in eukaryotic proteins, many predicted residues fall outside this high confidence regime.

Besides pLDDT, packing relationships have been recognized as a key feature for identifying regions of interest in AlphaFold predictions. Critical Assessment of protein Intrinsic Disorder (CAID) makes use of the AlphaFold_Bind measure as one of its baselines for identifying intrinsically disordered regions (IDRs) with conditional folding (Conte et al., 2023), and AlphaFold_Bind depends in part on accessible surface area (Piovesan et al., 2022). The tool AlphaCutter has been developed as an alternative method for preparing AlphaFold predictions for downstream structural biology uses, and it uses contact packing to identify and preserve folded regions with potential predictive value even at low pLDDT (Tam & Iwasaki, 2023).

We previously reported on the presence of such well-folded, protein-like regions within low-pLDDT (pLDDT < 70) predictions, calling them “near-folded” in reference to their presumed closeness to the target structure (Richardson et al., 2023). Other work has supported the predictive value of selected low-pLDDT regions, with residues having pLDDT as low as 40 being useful in constructing molecular replacement targets (Wang et al., 2025).

Selective identification of predictive or near-predictive residues is important because low-pLDDT regions are dominated by non-predictive residues, most obviously the *barbed wire* regions which appear highly disordered and hugely enriched in validation outliers. These regions must be removed for many structural biology tasks, including when preparing molecular replacement targets.

The relationship of AlphaFold predictions to disorder has been an area of considerable interest (Conte et al., 2023) (Necci et al., 2021) (Wilson et al., 2022). Unsurprisingly, low-pLDDT regions correspond strongly with intrinsically disordered regions (Tunyasuvunakool et al., 2021). However, high-pLDDT residues can also correspond with intrinsic disorder, especially regions of conditional folding (Alderson et al., 2023). pLDDT alone is not sufficient to identify modes of disorder within AlphaFold2 predictions.

This work formally categorizes different behaviors in AlphaFold2 predictions, especially the low-pLDDT regions, adding additional modes to our previous classification to make better sense of the ambiguous mid-confidence range from pLDDT ∼40-70. We also explore the relationships between our modes and protein disorder through comparison to disorder annotations from the MobiDB database (Piovesan et al., 2025). We present a tool to automatically split an AlphaFold prediction into our modes. This tool expands on AlphaCutter’s approach by adding MolProbity validation metrics (Williams et al., 2018) in addition to our own version of packing analysis (Davis et al., 2007). Our tool is written in Python and included in the Phenix software package (Liebschner et al., 2019) as `phenix.barbed_wire_analysis`, and in the Computational Crystallography Toolbox (cctbx) as `molprobity.barbed_wire_analysis`.

### 1. 2. Results

We surveyed low-pLDDT regions of AlphaFold predictions, using the human proteome predictions available from the AlphaFold Protein Structure Database (Varadi et al., 2024) as our dataset. The initial survey was visual, using MolProbity validation markup and all-atom contacts to identify patterns of behavior.

At low pLDDT, we observed three primary modes and one minor mode of AlphaFold prediction – all mutually exclusive – based on the combinations of high or low packing contacts with high or low density of validation outliers (Figure 1). At one extreme is the *near-predictive* mode, which strongly resembles folded protein. At the other is *barbed wire*, which has essentially no protein-like properties. By applying packing criteria to high-pLDDT regions, we defined two additional prediction modes, *predictive* and *unpacked high-pLDDT*, for a total of 6 modes.

**Figure 1.**
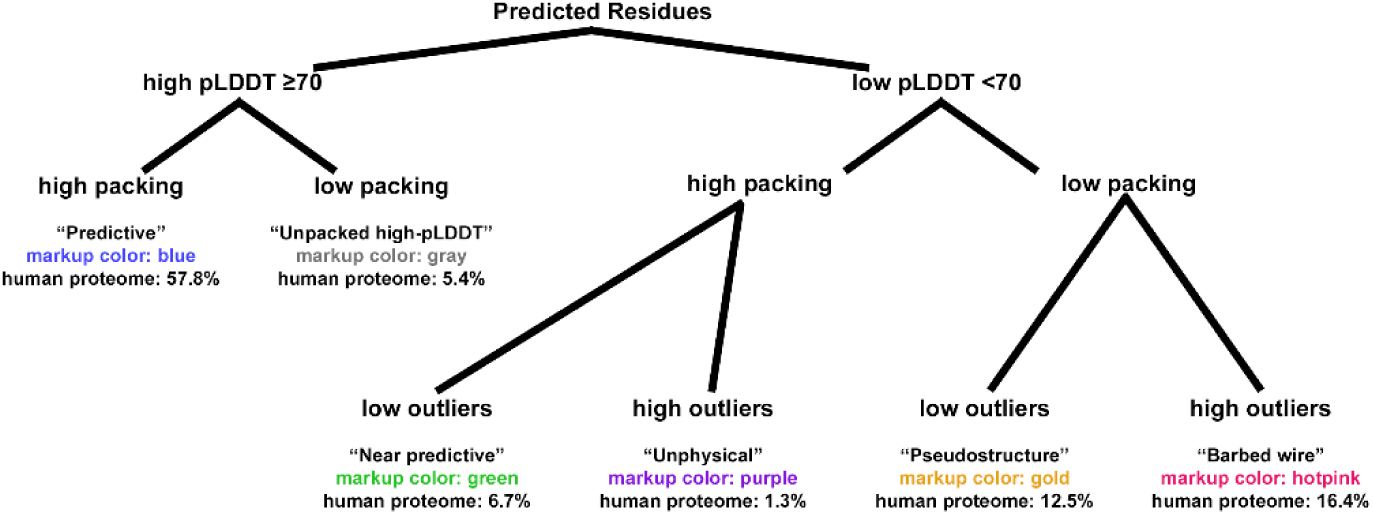
Prediction modes of AlphaFold2 and their relationships. This tree diagram shows how AlphaFold2 residues are divided into our modes, first by pLDDT, then by contact packing, then by validation outliers if necessary. Validation outliers are rare in high-pLDDT regions and are not used to define additional modes. For each mode, the name is listed, followed by the color used in our tool’s kinemage markup, followed by the frequency of that mode within the human proteome predictions. In the kinemage, markup for each mode can be toggled individually to aid readability. Example markup is shown in Figure 4.

We developed an analysis tool to categorize AlphaFold2 residues into these modes based on pLDDT, packing, and validation criteria. The tool can output text or JSON annotations of residues, a structure file pruned to include only residues of selected modes, or visual annotations in the form of kinemage markup viewable in our KiNG software (Chen et al., 2009). The markup color codes for each mode are shown in Figure 1, and the characteristics of each mode are described below.

### 2.1. Barbed wire

Many low-pLDDT regions are typified by wide, looping coils and by spike-like near-parallel arrangements of backbone carbonyl oxygens (Figure 2A). To our eyes, these features are reminiscent of coils of barbed wire (Figure 2B), so we have given them that name. “Spaghetti” is a metaphor in common use for these regions (Jones & Thornton, 2022, Varadi et al., 2024), but we do not favor the pasta analogy as it implies both flexibility (in contrast to the wide, rigid-looking arcs we observe) and closely piled packing (in contrast to the extremely low packing density we use to define this mode). Occasionally, these regions are referred to as “ribbon-like” (Abramson et al., 2024) (Piovesan et al., 2022), but we strongly disfavor that metaphor, given our historical connection to the development of ribbon diagrams for representation of protein alpha helices and beta sheets (Richardson, 2000), which *barbed wire* does not resemble.

**Figure 2.**
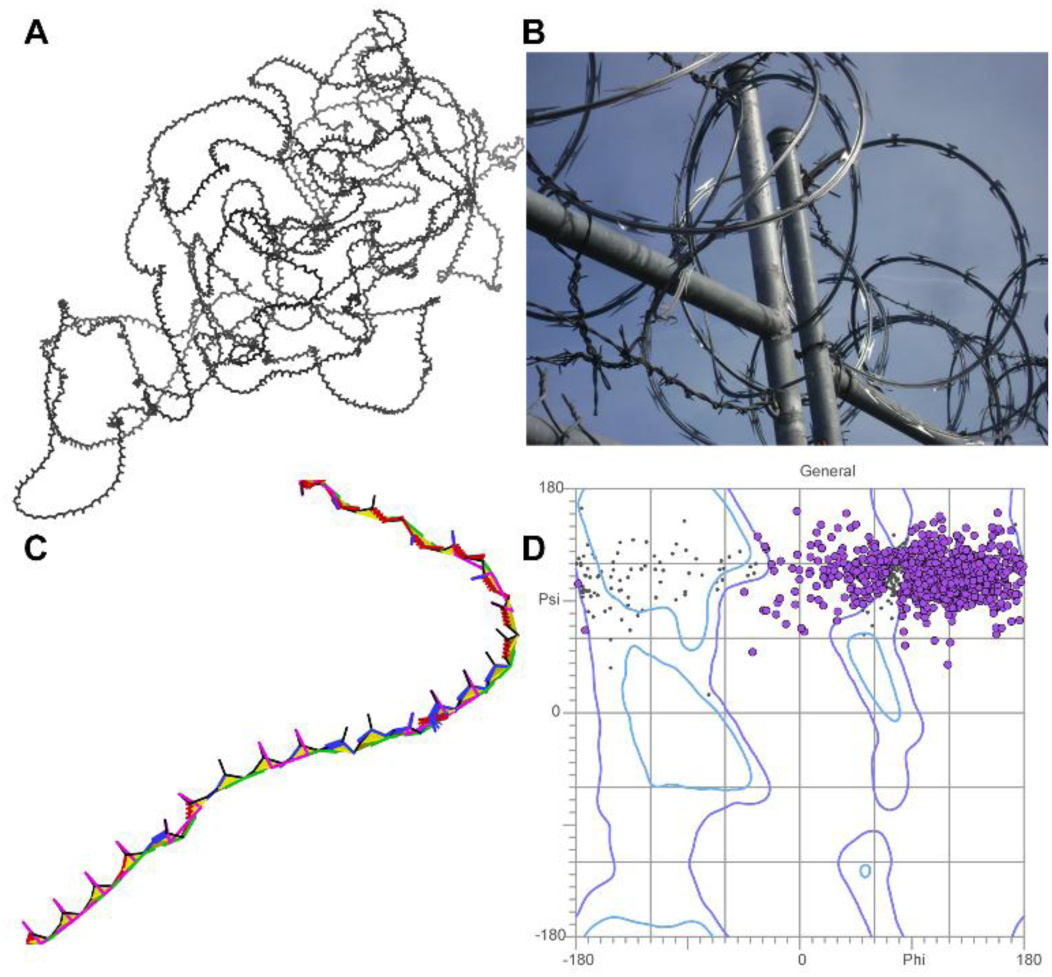
*Barbed wire* residues in Alphafold2 predictions. A: Nearly all-*barbed wire* prediction of fragment 6 of UniProt Q86YZ3 (very long sequences are predicted as multiple overlapping fragments). Wide, looping, or tangled coils are typical of the barbed wire prediction mode. B: Real barbed wire, whose spikes and coils give the prediction mode its name. (Image credit: Smithers7 by way of Wikimedia Commons, Creative Commons Attribution 3.0 Unported) C: Zoomed-in view of the Q86YZ3 fragment 6 prediction, residues 962-1005, with MolProbity validation markup, showing extremely high density of validation outliers. Markup is green for Ramachandran outliers, red and blue for covalent geometry outliers, magenta for CaBLAM, lime green and yellow for *cis* and twisted peptide bonds. CA geometry outliers from CaBLAM are omitted for clarity but are pervasive.

More diagnostic than these visual features is the extreme un-protein-likeness of *barbed wire* regions. *Barbed wire* residues are almost entirely unpacked – those great looping coils afford essentially no local steric contacts, and the coils generally do not pass near other parts of the structure. Additionally, *barbed wire* residues have an extremely high rate of backbone geometry outliers when assessed with MolProbity structure validation (Prisant et al., 2020). We observed that each residue in a *barbed wire* region typically manifests at least two (and frequently more) of the following: Ramachandran outliers, CaBLAM outliers, *cis* or twisted peptide bonds, covalent bond length outliers, and covalent bond angle outliers (Figure 2C). Ramachandran outliers in *barbed wire* regions occur primarily in the upper right portion of the Ramachandran plot (Figure 2D).

Cbeta deviation outliers (Lovell et al., 2003) are also common, though less pervasive or diagnostic. Surprisingly, given the density of other outliers, *barbed wire* residues show very few sidechain rotamer outliers or steric clashes.

Backbone covalent bond angles are of particular interest for diagnosing and understanding *barbed wire*. Although any of the backbone covalent bond angles within *barbed wire* may be distorted, we found that the C-N-CA bond angle is systematically abnormal in low pLDDT regions with high outlier density (Figure 3). The peak of the distribution for these C-N-CA angles falls at about -4σ, our typical cutoff for identifying bond angle outliers.

**Figure 3.**
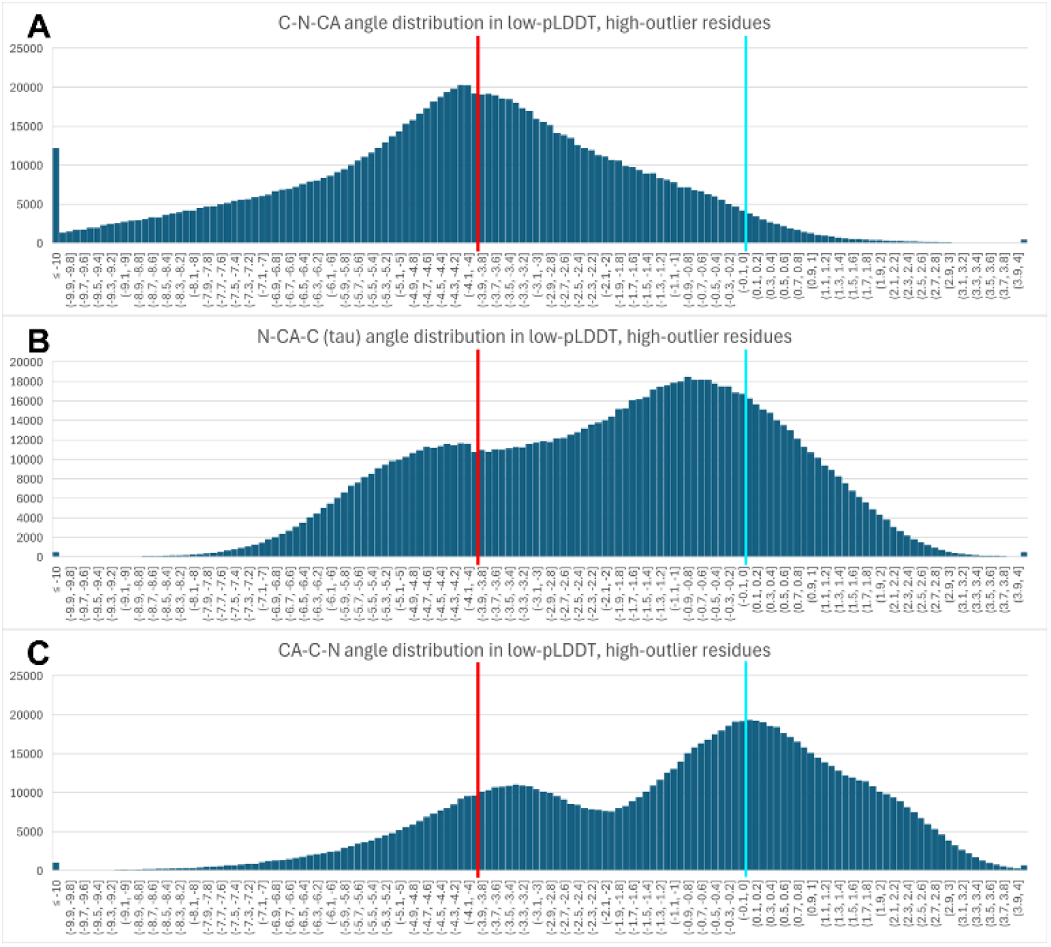
Histogram distributions of backbone covalent bond angles from AlphaFold2 prediction residues with low pLDDT, low packing, and high outlier density (these residues are *barbed wire*-like, but without explicit selection for angle outliers). The x axis is σ from target angle value, with bins of 0.1σ. The y axis is count of residues falling in the bin. Target (0σ) is marked with light blue bar; outlier threshold (−4σ) is marked with red bar. Underflow bin is ≤-10σ; overflow bin is >+4σ (the other outlier threshold). All three N/CA/C bond angles show frequent geometric distortions, with the C-N-CA angle’s distortion being systematic, and its distribution recentered almost exactly on the -4σ outlier threshold (A). This angle partially spans the peptide bond, and its systematic distortion suggests errors in peptide bond assembly.

Carbonyl oxygen bonds are frequently pointed in the same direction, rather than alternating as in beta strands. D: Ramachandran distribution for general-case residues in the Q86YZ3 fragment 6 prediction. Outliers are marked in purple. The distribution is highly unusual and clustered in the upper right of the plot, corresponding to an extended but unproteinlike conformation.

It is likely possible to define a geometric description for our visual intuition that *barbed wire* occurs in wide, rigid coils. However, this additional description was not necessary at this time. We found that the combination of packing and validation was sufficiently diagnostic, especially since C-N-CA angle outliers, *cis* and twisted peptide bonds, and upper-right quadrant Ramachandran outliers serve as consistent signatures for the *barbed wire* mode. (See Methods for a complete description of how signature outliers are treated.)

Users evaluating AlphaFold predictions with MolProbity or other validation software must take care. The presence of *barbed wire* regions will greatly worsen the apparent quality of whole-model validation statistics (e.g. overall Ramachandran outlier percent), without reflecting on the actual quality of the high-pLDDT or otherwise well-predicted regions of that structure. As always with validation, local details and context matter more than whole-model averages.

### 2.2. Near-predictive

At the other extreme from the underpacked and un-protein-like *barbed wire* are *near-predictive* regions. These are regions of low-pLDDT prediction that nevertheless have protein-like packing and geometry. At least some of these *near-predictive* regions are cases where AlphaFold has produced a mostly correct prediction, but has undervalued the confidence, resulting in a score below the critical pLDDT 70 threshold. *Near-predictive* regions can have a few validation outliers and steric clashes, similar to high-confidence regions and experimental structures, but do not have a diagnostic pattern of outliers.

After identifying *near-predictive* regions of human AlphaFold2 predictions, we surveyed the PDB in search of experimentally solved versions of those same regions to assess the accuracy of the predictions. This was largely an exercise in frustration – sequences that fold stably and behave well experimentally appear strongly correlated with sequences that AlphaFold2 predicts with high confidence. Finding experimental versions of *near-predictive* regions is rare because most residues deposited in the PDB have high-pLDDT AlphaFold counterparts. Frequently, *near-predictive* regions were not modeled in the PDB structures with corresponding UniProt IDs. Predictions such as UniProt Q9UM47 (Neurogenic locus notch homolog protein 3) where the *near-predictive* regions reflect a large number of repeated domains that are not present in the experimental structures were a systematic case of this problem. *Near-predictive* regions that are part of a small zinc finger motif, as in the prediction for UniProt O95218, were also common.

The best examples we found of *near-predictive* AlphaFold regions matching experimental structures are domains from eukaryotic translation initiation factor 3. The predictions of UniProt P60228 (Figure 4 A) and Q7L2H7 (Figure 4 D) have low and very low pLDDT confidence scores, respectively, but nevertheless align very well with chains E and M of PDB structure 6zon, experimentally solved at 3.0Å by electron microscopy (Thoms et al., 2020). There are some distortions, especially in the loops marked with purple spheres in Figure 4 E, which is part of why we designate this mode as *near-predictive* rather than fully-predictive. However, these predictions are clearly near enough to the target to be useful for many structural biology purposes, especially if supplemented with experimental data as in Phenix’s iterative Predict and Build pipeline (Terwilliger et al., 2023).

**Figure 4.**
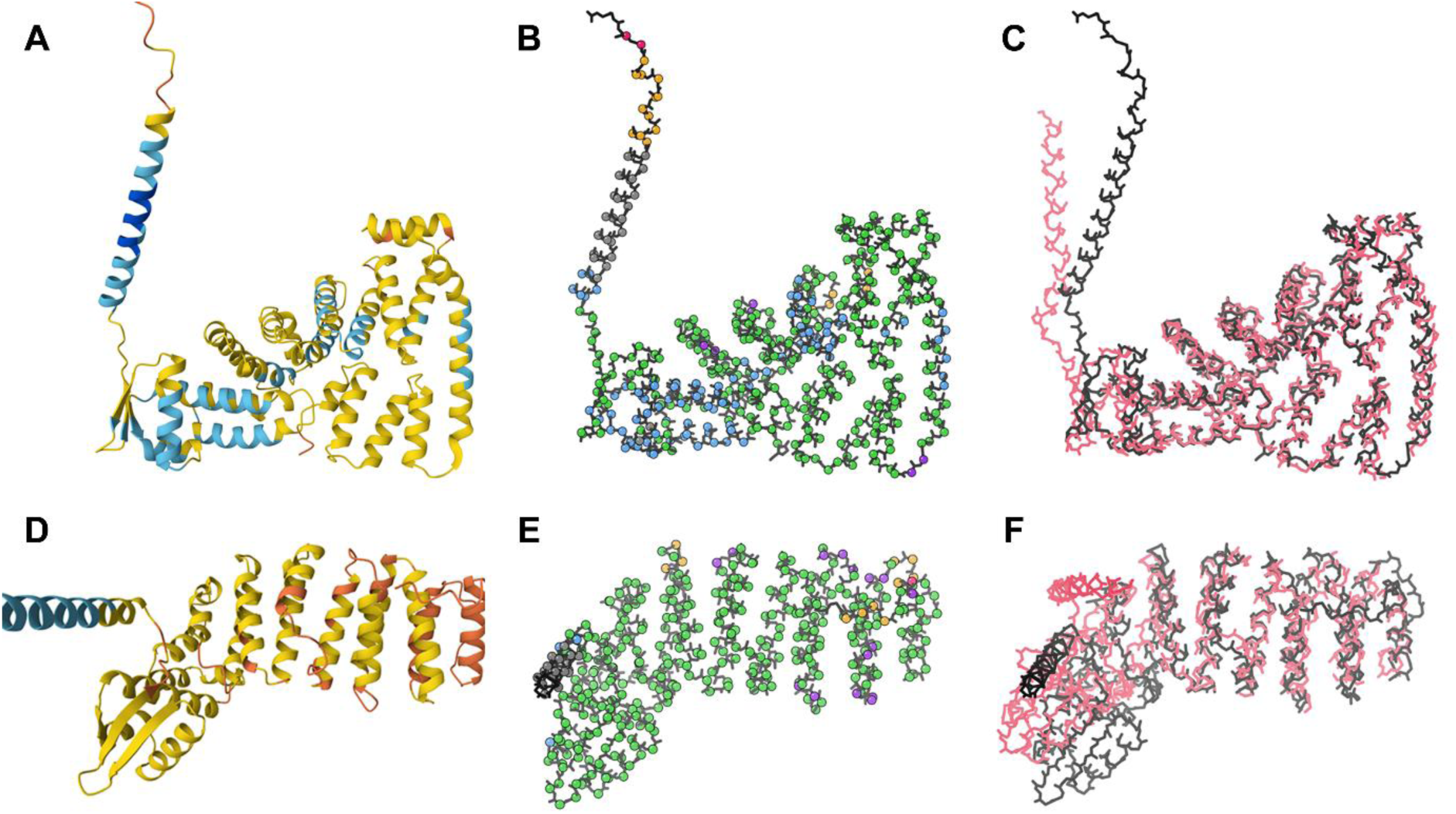
*Near-predictive* residues from AlphaFold2 predictions of eukaryotic translation initiation factor 3 and their experimentally-solved counterparts. A: Prediction of UniProt P60228 with standard pLDDT coloring, from the Mol* viewer at the AlphaFold Database. B: Prediction of UniProt P60228 with markup from our `barbed_wire_analysis` tool. The structure is primarily *predictive* (blue) and *near-predictive* (green), with the top left helix an example of *unpacked high pLDDT* (gray). C: Superposition of the prediction (black) with 6zon chain E (pink), solved by CryoEM at 3.0Å. D: Prediction of UniProt Q7L2H7 with standard pLDDT coloring. This prediction is lower confidence than A. E: Prediction of UniProt Q7L2H7 with markup from our `barbed_wire_analysis` tool. The structure is primarily *near-predictive* (green), with some *unphysical* (purple) regions. F: Superposition of the prediction (black) with 6zon chain M (pink). Even at low confidence, these predictions are close enough to the experimental structures to have structural meaning. The regions marked as *unphysical* – indicating severe validation outliers – correlate with loops omitted from the experimental structures.

There is some indication that AlphaFold3 (Abramson et al., 2024) more accurately assesses the confidence of regions that were *near-predictive* under AlphaFold2 and assigns them higher pLDDT scores. However, without a proteome-level resource of AlphaFold3 predictions, systematic study of AlphaFold3’s behavior in former and current low-pLDDT regions will be limited.

### 2.3. Pseudostructure

Between *barbed wire* and *near-predictive*, we observe a third mode of behavior, having protein-like residue geometry but minimal packing contacts. In addition to having legal peptide bond arrangements, *pseudostructure* residues generally assume conformations sufficiently similar to recognizable secondary structure elements (Wilson et al., 2022) that the Mol* viewer (Sehnal et al., 2021) at the AlphaFold Protein Structure Database draws many of them as ribbons instead of coil (Figure 5 A). However, on closer inspection, *pseudostructure* regions generally lack features of well-formed structure.

**Figure 5.**
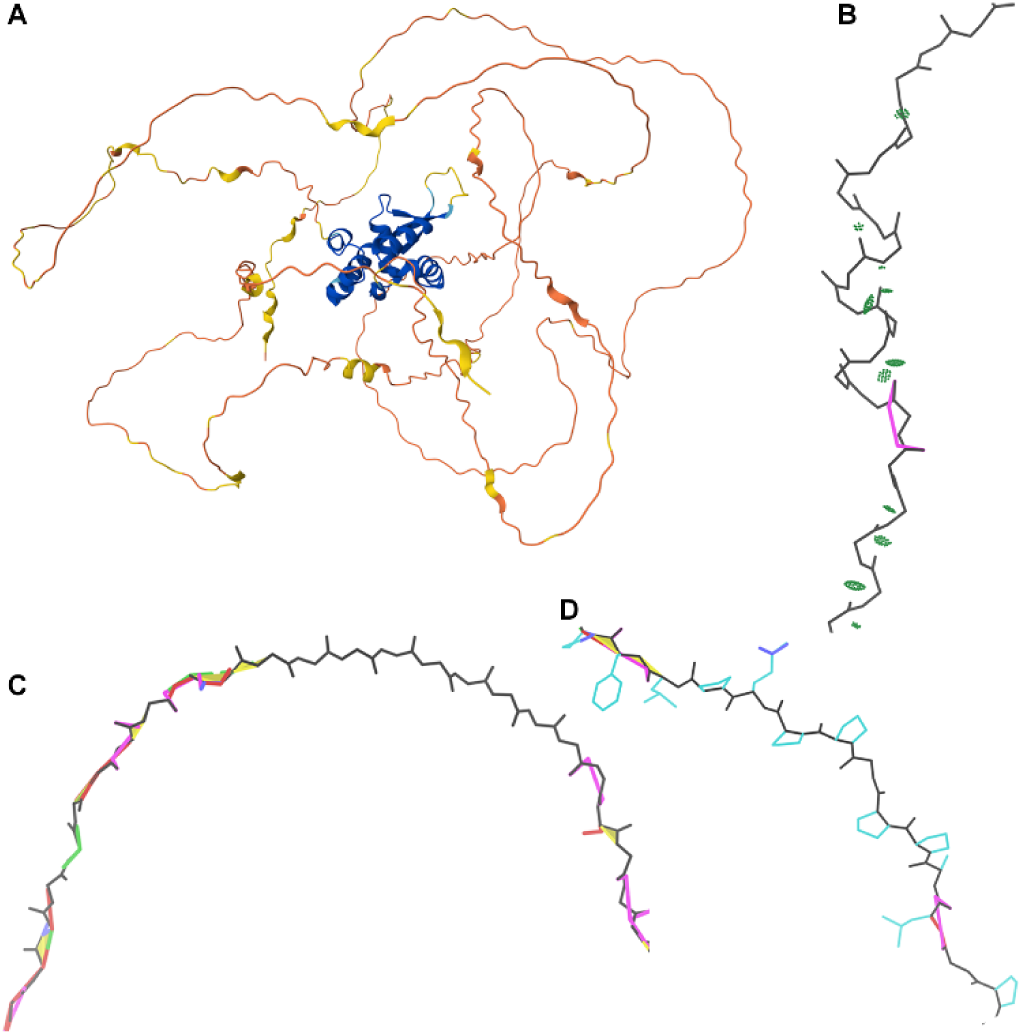
Examples of low-pLDDT *pseudostructure* predictions from UniProt O15353, human Forkhead box protein N1. A: Overall prediction colored by pLDDT, from the Mol* viewer at the AlphaFold Database. A well-packed and well-predicted core is surrounded by *barbed wire* and *pseudostructure*. Several *pseudostructure* elements are sufficiently similar to secondary structure to be depicted as ribbons by Mol*. B: A pseudostructure helix, residues 544-554. Hydrogen bonding (light green pillows) is inconsistent or weak, and the helix is not well-formed. A gamma-turn-like segment, a rare *pseudostructure* feature, is visible before/below the helix. C: A *pseudostructure* beta strand, residues 161-169. This strand is unpaired, so has no hydrogen bonding. *Barbed wire* regions before and after the strand show the sharp difference in validation outlier density, even though all of this strand is predicted at very low confidence (mostly pLDDT < 40). D: A poly-proline II region, residues 464-470. This conformation is correctly associated with regions of high proline content in AlphaFold predictions, but often occurs with low pLDDT.

*Pseudostructure* helices are often stretched or extended, and they have severely weakened or absent i-to-i+4 hydrogen bonding (Figure 5 B). *Pseudostructure* beta strands similarly lack hydrogen bonding, being isolated and unpaired (Figure 5 C). Geometry distortions in these beta strands are less obvious if present, as beta structure permits flexibility. However, some *pseudostructure* regions contain multiple Ramachandran outliers that populate a shoal just below and to the right of the general beta regions. This Ramachandran conformation is distinct from the typical *barbed wire* conformation, having negative rather than positive φ, and may represent a common distortion of beta-like structure at low pLDDT.

*Pseudostructure* also contains poly-proline II-like regions (Figure 5 D). Poly-proline II (Adzhubei et al., 2013) (Sasisekharan, 1959) is a repeating conformation formed by successive prolines with Ramachandran conformation around φ -75°/ψ 150°. Regions that AlphaFold2 predicts as poly-proline-like in conformation are indeed usually poly-proline-like in sequence, having a very high percentage of proline residues. These poly-proline regions may extend for more residues than poly-proline typically does in ordered regions of experimental structure, but are otherwise among the most consistently well-formed of the *pseudostructures*.

We also observe occasional regions of extended gamma-turn conformation (Matthews, 1972), with an i-to-i+2 hydrogen bonding pattern (Figure 5 B). This is a rare *pseudostructure* conformation and one of the only ways we observed to generate a consistent hydrogen bonding pattern at low pLDDT.

### 2.4. Unphysical

We find that high-pLDDT, predictive regions of AlphaFold2 structures are overall similar to well-solved experimental structures in MolProbity validation quality. Validation outliers occur at a reasonable rate (i.e. rarely, but not never) and are generally not diagnostic of some underlying prediction phenomenon. This behavior extends to *near-predictive* regions, which likewise generally conform to validation expectations.

Regions with high rates of both validation outliers and packing – designated *unphysical* – are therefore surprising. While *barbed wire* loops are generally well-separated from other structure in space, they occasionally pass close enough to other structure to appear packed. The most drastic form of this close approach is an actual intersection between parts of the chain, which results in strong steric clashes in addition to other severe validation outliers. Chain intersections were the only case where we systematically observed severe steric clashes at low pLDDT.

While most of these regions are properly understood as a variation on *barbed wire*, we give high-outlier high-packing regions the separate *unphysical* designation to draw attention to their association with impossible chain intersections.

Recognition of the *unphysical* mode is an advancement over AlphaCutter (Tam & Iwasaki, 2023). In our testing with its default settings, AlphaCutter tends to accept chain intersections and close approaches as valid globular-like structure due to their high contact packing and despite their physical impossibility.

An interesting case of *unphysical* residues can be seen in the Q7L2H7 eukaryotic translation initiation factor 3 subunit M prediction (Figure 4 E). The residues identified as *unphysical* (purple spheres) are largely at or near loops that have been omitted from the 6zon reference structure (Figure 4 F, pink backbone). These loops are short, and in the predicted model they do not extend far enough from the rest of the structure to lose packing. Validation outliers in the predicted model thus correspond to regions of greater flexibility or uncertainty in the experimental structure, and the *unphysical* designation marks residues with low predictive value.

### 2.5. High-pLDDT prediction modes

Contact analysis also elucidates an additional minor prediction mode at high pLDDT. Besides the main predictive mode, which is protein-like in its high degree of contact packing, there are underpacked regions nevertheless predicted at high or very high confidence. We call these unpacked high-pLDDT. Unpacked high-pLDDT regions are most often long helices that stick out from the well-predicted core of a structure (Figure 4 A). These regions will generally be explicable in the context of the structure, for example many are conditionally ordered (Alderson et al., 2023), or are membrane or domain insertion helices. All these cases may need to be trimmed from an experimental construct to facilitate crystallization.

### 2.6. pLDDT and prediction modes

We determined the pLDDT distributions of residues from the low-pLDDT prediction modes (Figure 6), excluding *unphysical* due to its rarity. pLDDT has a minimum observed value around 20. *Barbed wire* shows a strong and almost diagnostic preference for very low pLDDT. *Near-predictive* prefers higher pLDDT. *Pseudostructure* populates the full low-pLDDT range from 20 to 70, with some preference for very low pLDDT. The lines for *barbed wire* and *near-predictive* cross near pLDDT 50, which corresponds to the yellow to orange transition in conventional AlphaFold pLDDT coloring (Figures 4D and 5A). Most *barbed wire* residues can be avoided with a pLDDT 50 cutoff, but *pseudostructure* cannot be distinguished from the other modes by pLDDT alone and requires other analyses.

**Figure 6.**
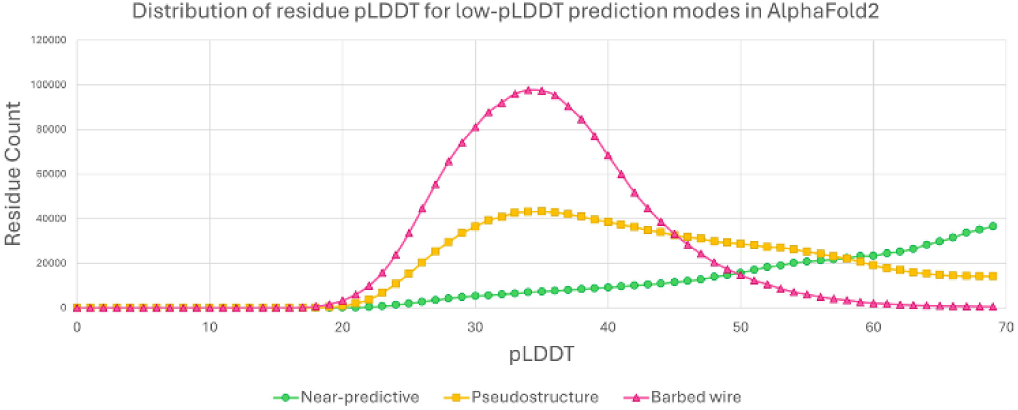
pLDDT distibutions for the major low-pLDDT prediction modes, pLDDT bins of 1, for sequences from the human proteome. *Barbed wire* (red, triangles) correlates with lower pLDDT scores. *Near-predictive* (green, circles) correlates with higher pLDDT scores. The crossing point for *barbed wire* and *near-predicitve* is close to 50, the yellow/orange boundary in conventional pLDDT coloring. If not for *pseudostructure* (gold, squares), which correlates only weakly with low pLDDT, a pLDDT 50 cutoff would be sufficient to select for most *near-predictive*.

The high and defined *barbed wire* peak supports our interpretation that *barbed wire* represents a distinct predictive behavior. The broad and weakly multi-modal distributions of *pseudostructure* and *near-predictive* suggest that these modes each contain multiple behaviors.

### 2.7. Sequence properties of prediction modes

The major prediction modes at low pLDDT – *barbed wire*, *pseudostructure*, and *near-predictive* – exhibit clearly different physical behaviors. We sought an explanation for why AlphaFold produces these different behaviors. In particular, is there a distinction between *barbed wire* and *pseudostructure* that might elucidate some property of disordered or partially ordered regions?

Anecdotally, AlphaFold2 predictions of nonsense sequences or text sequences seem to be dominated by the *pseudostructure* mode, rather than *barbed wire*. This suggests that mere nonsense is not sufficient to produce a *barbed wire* prediction. However, a thorough exploration of nonsense/random sequence prediction is beyond the scope of this work.

The MobiDB database (Piovesan et al., 2025) collects and presents many annotations and predictions of disorder. We surveyed regions of *barbed wire*, *pseudostructure*, *near-predictive*, and high-pLDDT AlphaFold2 prediction modes for correlations with MobiDB annotations. To reduce the effects of edge definitions and smoothing, we only considered residues from uninterrupted (pre-smoothing) segments of *barbed wire*, *pseudostructure*, and *near-predictive* that were at least three residues long after the first and last three residues were removed from the segment. This aggressive pruning focused the survey on only unambiguous cases of each prediction mode. High-pLDDT regions were identified with a simple pLDDT >= 70 cutoff and include both our *predictive* and *unpacked high-pLDDT modes*.

The complete MobiDB frequency analysis is provided in the Supporting Information; Figure 7 excerpts points of interest from the complete analysis. Many of the MobiDB annotations show a stair-step pattern, with high-pLDDT residues the least correlated with disorder annotations, then *near-predictive*, then *pseudostructure*, and then *barbed wire* being the most correlated with disorder (Figure 7, prediction-disorder-iupl). This general pattern suggests that the difference between *barbed wire* and *pseudostructure* is one of degree rather than type.

**Figure 7.**
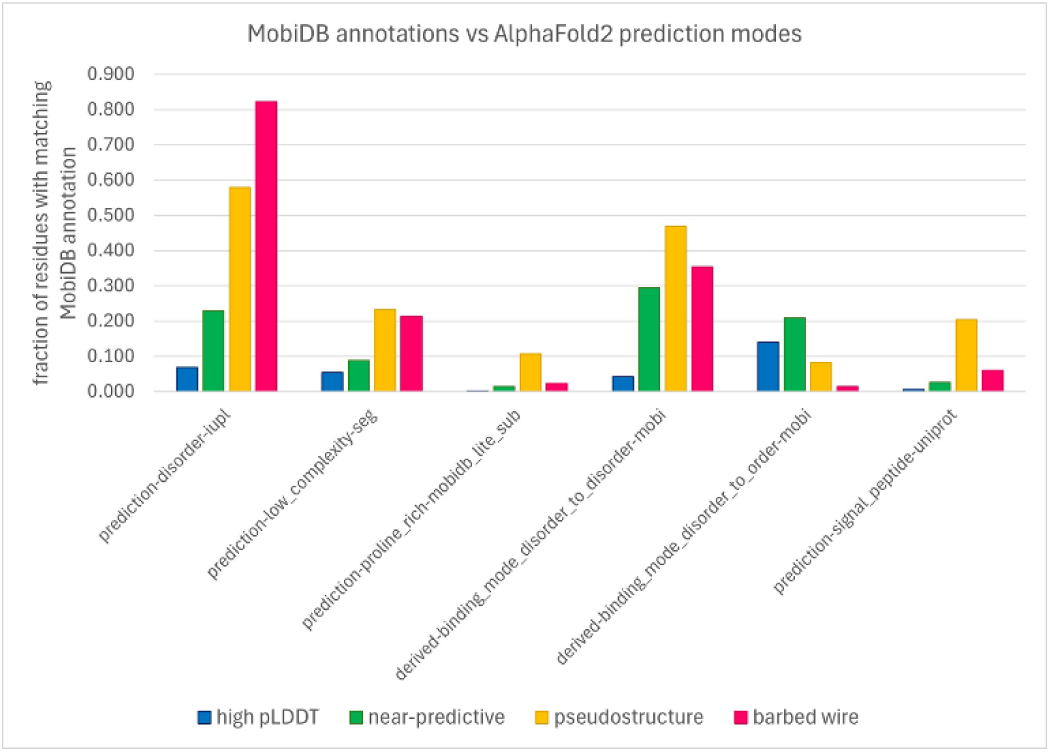
Relationships of AlphaFold2 prediction modes with MobiDB disorder annotations. From left to right: prediction-disorder-iupl shows a typical distribution for many predictions of disorder (see Supplement), where high pLDDT residues are least associated with disorder and *barbed wire* residues are most associated, but no mode stands out as exceptional. The distributions to the right are exceptions to this general pattern. Low complexity sequences are about equally associated with both *pseudostructure* and *barbed wire*. Proline-rich sequences are preferentially associated with *pseudostructure*, where we observe the poly-proline II conformation. Disorder-to-disorder binding somewhat favors *pseudostructure*. Disorder-to-order binding somewhat favors *near-predictive*. Predicted signal peptides strongly favor *pseudostructure* over any other mode.

However, certain annotations break from this pattern. We had hoped that low-complexity sequences would explain our prediction modes. The prediction-low_complexity-seg annotation indeed shows higher preference for the non-predictive modes than for either of the predictive modes, but unfortunately does not distinguish between the non-predictive modes. In retrospect, we recognize that low-complexity sequences are diverse in behavior and not monolithically associated with either order or disorder. For example, prediction-proline_rich-mobidb_lite_sub shows strong preference for the *pseudostructure* mode over any other, in agreement with our observation that poly-proline-II conformations are regularly identified as *pseudostructure* in our system.

Derived-binding_mode_disorder_to_disorder-mobi shows some preference for *pseudostructure* over the other low-pLDDT modes. Derived-binding_mode_disorder_to_order-mobi shows a preference for *near-predictive* and a strong preference against *barbed wire*. The low pLDDT of *near-predictive* regions may therefore reflect a correlation with regions of conditional rather than permanent order. These results suggest that regions AlphaFold2 predicts as *barbed wire* are not correlated with even conditional order. Annotations from IDEAL (Fukuchi et al., 2012), a method focused on regions of conditional order, show similar patterns (see Supporting Information Figures S4 and S5).

The most interesting single preference revealed by the MobiDB survey is prediction-signal_peptide-uniprot. This annotation shows strong preference for *pseudostructure* above any other mode. This may represent a case where AlphaFold2 can recognize, by chance or design, a specific sequence property relevant to structural biology outside of folded domains.

## 1. 3. Methods

### 3.1. Dataset

We downloaded the complete human proteome prediction from the AlphaFold Protein Structure Database (Varadi et al., 2024) (version 2 initially, version 4 when it became available – the differences in these versions did not appear to impact our analysis). We chose the human proteome because human proteins contain complex binding, regulation, and generally extensive intrinsically disordered regions, making them challenging cases for AlphaFold and good test cases for examining low-pLDDT predictions, especially *barbed wire*.

The human proteome download contains predictions for some very long protein sequences, which were broken into overlapping fragments for prediction. Predictions treated in this fashion were useful in the early stages of our survey, as they provided examples with especially high *barbed wire* content (Figure 2). Fragmented predictions were excluded from our final analyses, as the fragmented sequences lacked their full context, the predictions appeared to be worse than for non-fragmented sequences, and the proper way to count the overlapping regions of the fragments was unclear.

### 3.2. MobiDB survey

For each AlphaFold2 structure we categorized, we downloaded the corresponding MobiDB entry based on UniProt ID, in JSON format. Each MobiDB entry provides ranges of residues from that sequence that fit within a variety of disorder categories. For each of our categorized residues, for each MobiDB disorder category, we determined whether or not the residue was within the MobiDB residue range, and added up the totals. Results are presented as the fraction of our categorized residues that share each MobiBD disorder annotation.

### 3.3. phenix.barbed_wire_analysis tool

The tool accepts a structure file in PDB or mmCIF format, with pLDDT in the B factor field, as per AlphaFold standard. Hydrogens are added to a submitted structure with Reduce, and contact analysis is performed with Probe (Davis et al., 2007). Secondary structure elements are identified based on CA geometry. For each residue, a packing score is determined based on the number of different steric contacts (0.25Å van der Waals surface separation or closer) per non-hydrogen atom in a 5-residue window (i-2 to i+2) around the residue of interest. Where secondary structure is identified, contacts internal to a given secondary structure element (such as the H-bonds between beta strands) are not counted. Likewise, local contacts within a sequence distance of 4 are omitted from the count. The purpose of these omissions is to focus on contacts from tertiary structure and ensure that helices and sheets must touch other elements of the structure to count as packed, despite their rich internal contacts. For helix and coil residues, a score of > 0.6 contacts per heavy atom is considered adequately packed. For beta strand residues, whose packing may be dominated by (omitted) intra-sheet contacts, the cutoff score is lowered to > 0.35.

MolProbity validations are run via Phenix (Liebschner et al., 2019), namely `ramalyze` for Ramachandran (Lovell et al., 2003) (Ramachandran et al., 1963), CaBLAM (Williams et al., 2018), `omegalyze` for cis and twisted peptide bonds (Williams et al., 2018), and `mp_validate_bonds` for covalent bond geometry (Moriarty et al., 2014).

A residue is marked as having a high density of outliers if two or more of the following are true about a three-residue window centered on that residue: two or more residues have *cis*-nonPro or twisted peptide bonds, two or more residues have CaBLAM and/or CA geometry outliers, two or more residues have covalent bond length and/or angle outliers, or all three residues fall in a high-psi Ramachandran band (+60 < ψ < +170) and at least one residue is a Ramachandran outlier.

An individual residue is marked as having a signature *barbed wire* outlier if it is a Ramachandran outlier falling in the upper right of the Ramachandran plot (−15 < φ < +170, +60 < ψ < +170). A residue is also marked as having a signature *barbed wire* outlier if it is a CA geometry outlier as determined by CaBLAM and is also in a low-pLDDT, unpacked region (this outlier is permitted in predictive regions.) Since *barbed-wire*-ness appears to be a property of joining residues together, both residues that share a peptide bond are marked as having a signature outlier if that peptide bond contains a C-N-CA bond angle outlier, a *cis*-nonPro peptide bond, or any twisted peptide bond. A *cis*-Pro is permitted in predictive regions and is only considered a signature outlier if it appears in a low-pLDDT and unpacked region.

Residues are then categorized into our defined modes (Figure 1). *Predictive* residues are high-pLDDT, high packing. *Unpacked high-pLDDT* residues are high-pLDDT, but low packing. *Near-predictive* residues are low-pLDDT, high packing, low outliers. *Pseudostructure* residues are low-pLDDT, low packing, low outliers. *Barbed wire* residues are low-pLDDT, low packing, and show high outlier density, a signature *barbed wire* outlier, or both. *Unphysical* residues are the relatively rare low-pLDDT, high packing, signature outlier/high outliers case.

To reduce fragmentation and simplify the annotation, isolated residues are smoothed into their neighbors. If 1 or 2 residues of one low-pLDDT category are surrounded on both sides by residues of another category, the surrounded residues are recategorized to match their surroundings.

Kinemage markup may be generated with `phenix.barbed_wire_analysis output.type=kin`. Besides color-coded balls on each CA (see Figure 1), the kinemage markup includes text label annotations for each residue showing how similar to *barbed wire* it is. L for low pLDDT, p for low packing, r for Ramachandran, o for omega (peptide bond dihedral), c for CaBLAM, g for covalent geometry. These labels always occur in the same order, Lprocg, and each may be replaced with a dash if the residue is not like *barbed wire* in that regard. Thus L is the typical label for *near-predictive* residues, and Lp is the typical label for *pseudostructure*.

A pLDDT 70 cutoff suffices for most AlphaFold prediction preparation. In cases where significant regions of a prediction are below pLDDT 70, the barbed wire analysis tool can provide an alternative preparation with `phenix.barbed_wire_analysis output.type=selection_file`, printing a PDB file that contains only residues from selected modes. By default, predictive and near-predictive residues are returned, but any combination of modes may be selected with a `modes=` flag. However, in challenging cases, we recommend leveraging the “intelligence augmentation” (Engelbart, 2023) of the complete kinemage markup rather than relying on automation.

## 1. 4. Discussion

We identify several mutually-exclusive modes of AlphaFold2 prediction behavior based on combinations of structure validation and packing criteria (Figure 1). These are *predictive* (high pLDDT, high packing), *unpacked high-pLDDT* (high pLDDT, low packing), *near-predictive* (low pLDDT, high packing, low outliers), *pseudostructure* (low pLDDT, low packing, low outliers), *barbed wire* (low pLDDT, low packing, high outliers), and the relatively rare *unphysical* (low pLDDT, high packing, high outliers).

While a simple pLDDT >= 70 cutoff is sufficient for selecting the good parts of a prediction in most cases, understanding these more detailed prediction modes can be valuable for properly using a prediction. For example, the *unpacked high-pLDDT* mode often contains insertion helices which may need to be truncated from a construct to achieve crystallization. More positively, the *near-predictive* mode contains residues which, even at low confidence, may be closely related to the real structure. *Near-predictive* residues would be prime candidates for an iterative prediction process such as Phenix’s `predict_and_build`.

Contact packing is the single most important criterion for identifying *near-predictive* regions at low pLDDT, since serious validation outliers within well-packed regions are rare. However, adding MolProbity validation criteria allows our *unphysical* category to annotate both residues of significantly worse prediction in otherwise *near-predictive* regions (see Figure 4 E) and sites of chain intersections, both of which can appear highly packed despite their errors. Validation criteria also introduce a possible distinction between *barbed wire* and *pseudostructure*.

### 4.1. Origins of prediction modes

Application of MolProbity validation reveals an apparent distinction between the *barbed wire* mode and the *pseudostructure* mode based on *barbed wire’s* overwhelming density of backbone geometry errors. This is supported by *pseudostructure’s* resemblance to secondary structure elements compared to *barbed wire’s* unproteinlike conformation.

We speculate that the typical *barbed wire* conformation and its attendant validation outliers are not the result of AlphaFold prediction, but rather the lack of a prediction. These are probably regions that are effectively unpredicted, where the residues are strung together according to a default behavior, to fill the necessary sequence space between predictable regions. This speculation that *barbed wire* represents a default behavior is supported by the strong association of validation outliers with the peptide bond. The commonness of C-N-CA bond angle outliers and *cis* and twisted peptide bonds indicates a consistently un-protein-like assembly of adjacent residues. The unusual distribution of *barbed wire* residues in Ramachandran space also suggests a default behavior. As shown in Figure 2D, *barbed wire* residues are typically distributed in a band at high ψ (+60 to +170) with a weaker preference for high φ (roughly -15 to +170). This implies that the starting value of the “residue gas” from which AlphaFold predictions are constructed is at ψ ≈ +110 (ψ is the only Ramachandran angle specified for a single non-proline residue, since the O_i_ position also defines the N_i+1_ position across the planar peptide bond), and that the deviations from that value are needed to prevent clashes.

If this speculation is correct, then the high frequency of validation outliers in *barbed wire* regions may be considered a positive feature of such AlphaFold predictions, rather than a failure. The combination of low pLDDT and high outliers clearly marks *barbed wire* as distinct from regions where AlphaFold has seriously attempted a prediction, most importantly *near-predictive*.

If *barbed wire* is an arbitrary assembly, then *pseudostructure* may be an attempted, but unsuccessful, prediction. The absence of validation outliers in *pseudostructure* suggests a more complete process. However, it is also possible that AlphaFold2 has multiple patterns of arbitrary assembly and that *pseudostructure* represents patterns that happen not to generate *barbed wire’s* signature outliers. If this is the case, then *pseudostructure* is not fundamentally different from *barbed wire*, and is likewise not an attempted prediction. Our current analysis cannot resolve the question of how – or whether – *pseudostructure* differs from *barbed wire*.

Poly-proline II presents a particularly ambiguous case. We observed above that the ψ Ramachandran dihedral is defined by the geometry within a residue and that therefore the distinctive Ramachandran distribution of *barbed wire* residues (Figure 2D) may reflect a default conformation within AlphaFold’s residue gas. In prolines, the φ dihedral is also heavily restricted by the sidechain geometry. The resulting φ,ψ combination would put a proline-rich residue gas in the correct Ramachandran region for poly-proline II. Thus, at least for proline, even an arbitrary assembly process might generate the geometrically legal and sequentially correct poly-proline II conformations we observe.

### 4.2. Sequence properties of prediction modes

Is there a sequence property that explains why AlphaFold produces such different-looking predictions in *pseudostructure* regions versus *barbed wire*? And if so, is that property relevant to structural biology, for example through correlation with intrinsically disordered regions (IDRs)? Our survey of disorder annotations from MobiDB was inconclusive. Broadly speaking, *near-predictive* regions had the most ordered-like residues at low pLDDT, according to MobiDB, and were substantially more ordered-like than *pseudostructure*. But *pseudostructure* was only somewhat more ordered-like than *barbed wire*, overall. Despite the association of signal peptides with *pseudostructure*, no MobiDB annotation category offered a clear explanation for why AlphaFold predicts some residues as *barbed wire* and others as *pseudostructure*. Nevertheless, we hope that this tool will be of use to others with a more nuanced understanding of disordered proteins.

AlphaFold2 is heavily dependent on multiple sequence alignment (MSA) for generating predictions. It is therefore possible that MSA depth could explain the barbed wire/pseudostructure distinction. The AlphaFold Protein Structure Database datasets do not report MSA depth information, so a complete analysis is not practical. However, MobiDB reports MSA occupancy for some sequences.

MSA occupancy is a per-residue score denoting the fraction of MSA sequences that contain a value for that residue. A high occupancy fraction indicates that most of the aligned sequences include that residue position. Only about 10% of the human proteome sequences from our study have MobiDB homology-msa_occupancy-psiblast annotations. Our analysis of MSA occupancy is therefore limited, especially since it is likely that the sequences which received homology-msa_occupancy-psiblast annotations are not independent of other sequence properties that might affect AlphaFold prediction. Figure 8 shows probability density histograms for our six AlphaFold2 prediction modes. The non-predictive modes – *barbed wire*, *pseudostructure*, and *unphysical* – show similar distributions to each other. *Barbed wire* has a lower high-occupancy peak relative to *pseudostructure* and a higher distribution across the low-occupancy range. As with many of the other MobiDB annotations discussed above, barbed wire shows somewhat greater correlation to disorder than *pseudostructure*, but not enough to provide a clear explanation.

**Figure 8.**
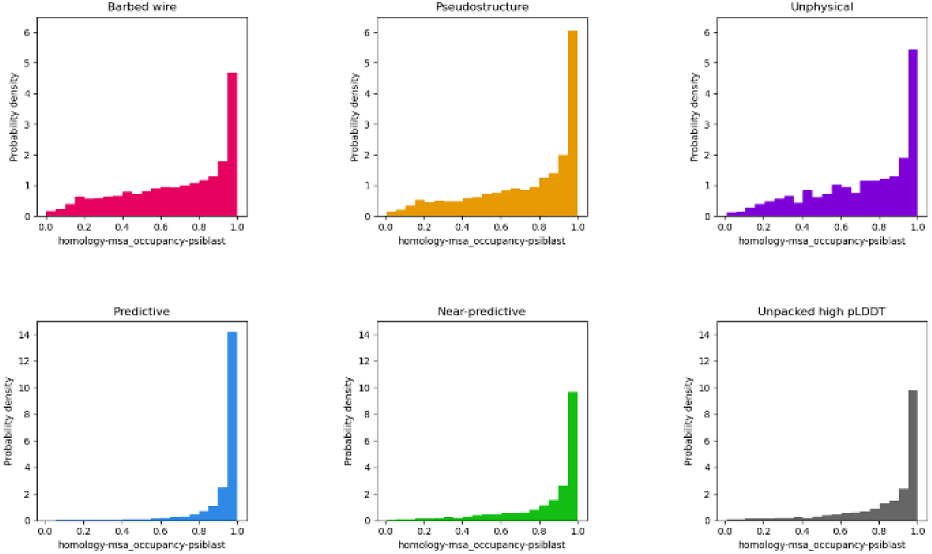
Multiple sequence alignment occupancy histogram distributions for AlphaFold2 human proteome predictions, as annotated in MobiDB. The three non-predictive modes (*barbed wire*, *pseudostructure*, and *unphysical*) show similar distributions to each other. *Near-predictive* has a distribution more similar to the high-pLDDT modes, supporting our association of *near-predictive* regions with *predictive*. Similarity between *near-predictive* and *unpacked high-pLDDT* suggests that *near-predictive* also contains conditionally binding regions.

*Near-predictive* has a distribution much more similar to the high-pLDDT modes (*predictive* and *unpacked high-pLDDT*) than to the other low-pLDDT modes. This similarity supports our assumption that *near-predictive* regions are generally near-correct predictions – they have MSA quality similar to *predictive* regions – but also reveals a more complex relationship with conditional folding.

### 4.3. Conditional folding

Work by Alderson et al. (Alderson et al., 2023), which focuses on relationships between intrinsic disorder and high-pLDDT regions, provides a useful companion piece to our work and is recommended. In particular, Alderson et al. explain many of the helices we identify as *unpacked high pLDDT* as regions of conditional order. They show that IDRs with high pLDDT (pLDDT ≥ 70) AlphaFold2 predictions have favorable MSA properties (high alignment depth and high conservation) relative to IDRs with low pLDDT (pLDDT < 50) predictions. However, they omit pLDDT between 50 and 70. Our tool can separate this difficult pLDDT range into major components of *near-predictive* and *pseudostructure* (Figure 6). We find that *pseudostructure* residues have unfavorable MSA quality similar to *barbed wire* (Figure 8), and therefore similar to residues predicted with pLDDT < 50. We find that *near-predictive* residues have favorable MSA quality similar to *unpacked high-pLDDT* residues, and therefore similar to the category of conditionally folding residues reported by Alderson et al. Our survey of MobiDB annotations also found that disorder-to-order binding was associated with high-pLDDT and *near-predictive* regions, but not with *pseudostructure* or *barbed wire* (Figure 7).

Taken together, these observations suggest that conditionally folded regions are an important component of AlphaFold2 predictions in the pLDDT 50-70 range, and that these regions are part of our near-predictive category. If *near-predictive* is populated by conditional folding (rather than consistent folding), that would help explain why experimentally-solved versions of *near-predictive* regions were difficult to find in the PDB. The packing-based *near-predictive* versus *pseudostructure* distinction is especially important for identifying potential conditional folding, since *barbed wire* can be largely removed via a pLDDT cutoff.

However, as the eukaryotic translation initiation factor 3 examples illustrate (Figure 4), the *near-predictive* mode is not exclusively conditionally folding. Much as the broader *pseudostructure* mode includes several distinct behaviors, *near-predictive* is not monolithic. Further investigation is needed to identify and interpret the most useful behaviors within this mode, and to understand how prediction of these regions changes from AlphaFold2 to AlphaFold3.

### 4.4. On templates

An important question, especially regarding *near-predictive* regions, is how much template structures from the PDB affect AlphaFold’s predictions. As with many questions about machine learning methods, a clear answer is difficult to extract. Anecdotally, it seems that where a good multiple sequence alignment exists, that MSA dominates in AlphaFold2, and presence or absence of templates matters less. However, it is also known that templates can be used to add positive information into AlphaFold during iterative prediction (Terwilliger et al., 2023). The apparent scarceness of experimental structures corresponding to *near-predictive* regions may suggest that where a PDB template is used, AlphaFold predicts with high confidence (this would be a generally desirable behavior). Or it may suggest that *near-predictive* residues correlate with regions that are also difficult to solve experimentally (e.g. conditionally folding).

### 4.5. On “Hallucinations”

We have intentionally avoided the term “hallucination” in describing low-pLLDT prediction modes. Hallucination implies predictive effort, so if *barbed wire* is indeed a non-predictive, default behavior, hallucination does not precisely apply. *Pseudostructure* might be understood as a hallucination, having some of the “grammar” of real structure, but little of the meaning. However, *pseudostructure* does not have sole claim to structural hallucinations within AlphaFold. Indeed, the most dangerous hallucinations are those that present plausible structures, either in our *near-predictive* mode or in high-pLDDT regions that have the wrong fold despite their confidence. Thus “hallucination” is a category distinct from but sometimes overlapping with the predictive modes described here.

## 1. 5. Conclusion

Development of structure prediction methods continues to advance. Alphafold3 appears to behave differently from AlphaFold2 in low-pLDDT regions (Abramson et al., 2024). AlphaFold3 may assign higher confidence to previous *near-predictive* regions and may be more ambitious in creating folded (possibly hallucinatory) structure in previous *barbed wire* regions. This work was made possible by the availability of proteome-level predictions for AlphaFold2. The above descriptions of low-pLDDT behaviors, especially the signature outliers, are therefore specific to AlphaFold2 and apply only partially to AlphaFold3 or other prediction systems (Baek et al., 2021) (Lin et al., 2023). However, the AlphaFold3 code was only recently released at the time of this writing, and AlphaFold3 is not broadly implemented. The Computed Structure Models served by the AlphaFold Protein Structure Database and the RCSB PDB are still AlphaFold2 models, and Phenix still uses AlphaFold2 in its integrated Predict and Build pipeline. Regardless of the advancements or changes in AlphaFold3, AlphaFold2 models remain a significant presence in structural biology.

In our work with MolProbity structure validation, we have always found visual inspection of models to be a vital part of understanding structures, especially in difficult cases – and AlphaFold predictions are by definition difficult cases because their means of production is obscured behind machine learning. The `phenix.barbed_wire_analysis` tool facilitates such an understanding of AlphaFold2 predictions.

## Supporting information

MobiDB annotations table

## Acknowledgements

The authors wish to thank Michael Prisant, who contributed to the literature search and critical reading of the manuscript. They also acknowledge Matt Baker, who shared a structure prediction containing a near-predictive region, which initially inspired this work.

## Conflicts of interest

The authors declare no conflicts of interest.

## Data availability

MobiDB survey results are available as an Excel file in the supporting information.

## Supporting information

**Figure S1.**
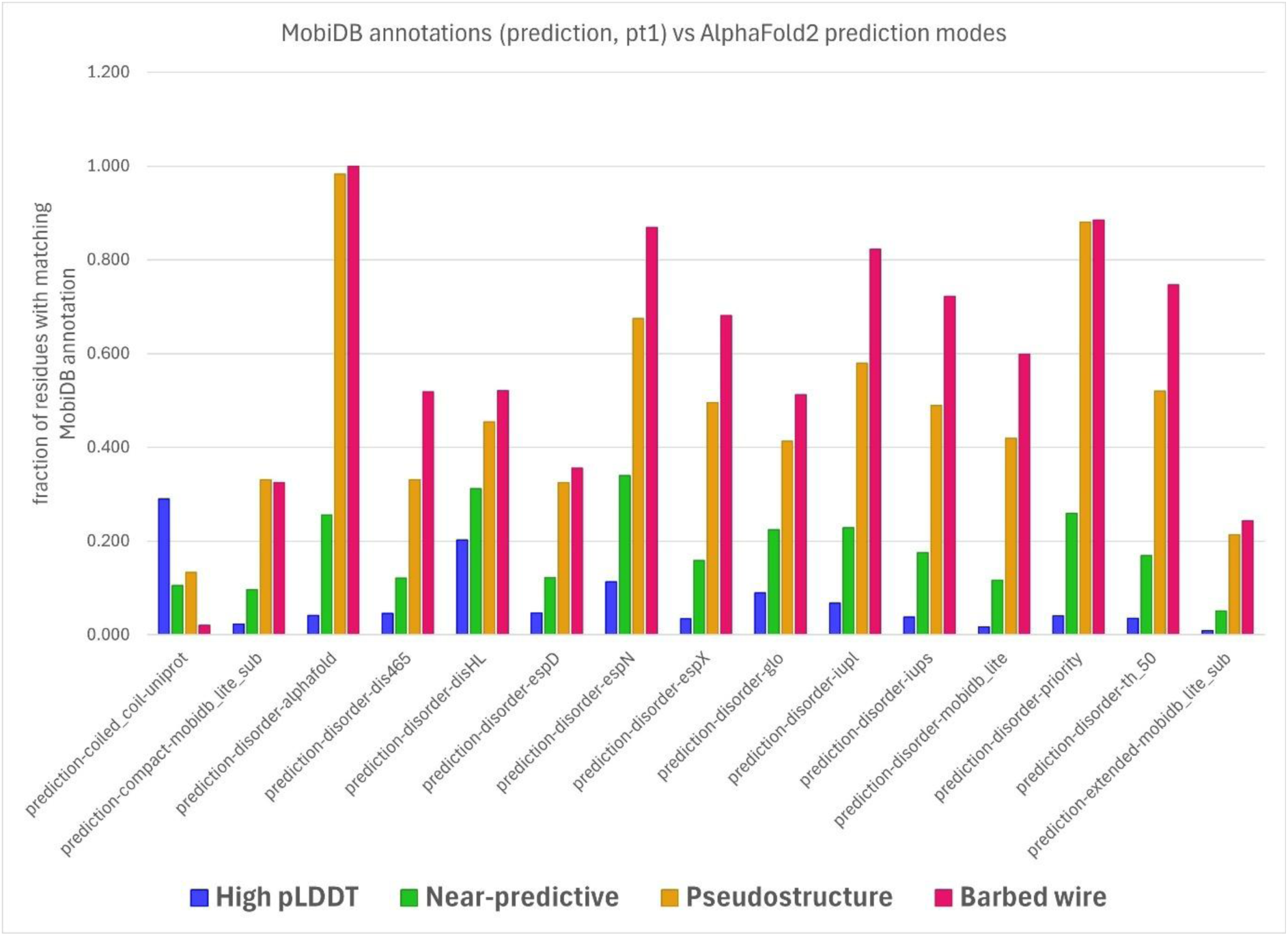
Prediction annotations from MobiDB and their relationships with AlphaFold2 prediction modes. Bar height is the fraction of residues from that prediction mode that were marked with the matching MobiDB annotation. Not all sequences are treated with all annotations, and only residues from duly annotated sequences were considered for each annotation. These prediction annotations show the general stair-step pattern discussed in the main text. Prediction-disorder-iupl is included in Figure 7 as a representative of this behavior. Correlations with prediction-disorder-alphafold are not considered significant, since that annotation is also interpreting AlphaFold results, rather than directly interpreting the underlying sequence.

**Figure S2.**
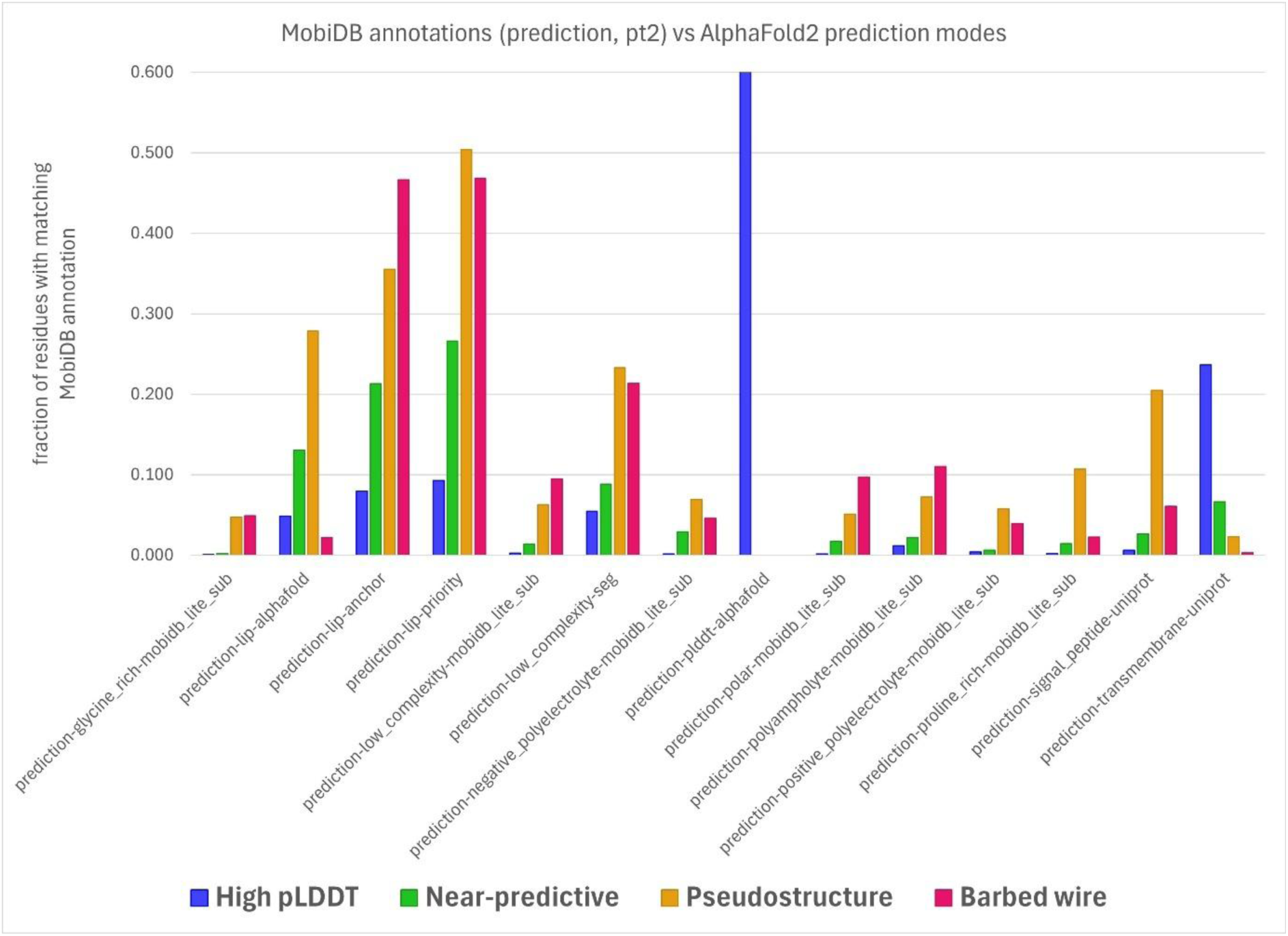
Additional prediction annotations from MobiDB and their relationships with AlphaFold2 prediction modes. Bar height is the fraction of residues from that prediction mode that were marked with the matching MobiDB annotation. Not all sequences are treated with all annotations, and only residues from duly annotated sequences were considered for each annotation. The y-axis is truncated at 0.6, otherwise the trivial prediction-plddt-alphafold result (which goes to 1.0) would dominate. Prediction-low_complexity-seg, prediction-proline_rich-mobidb_lite_sub, and prediction-signal_peptide-uniprot are included in Figure 7. The preference of prediction-transmembrane-uniprot for high-pLDDT residues partly reflects the tendency of membrane insertion helices to be predicted in the *unpacked high-pLDDT* mode, which is included in the high-pLDDT category for these plots.

**Figure S3.**
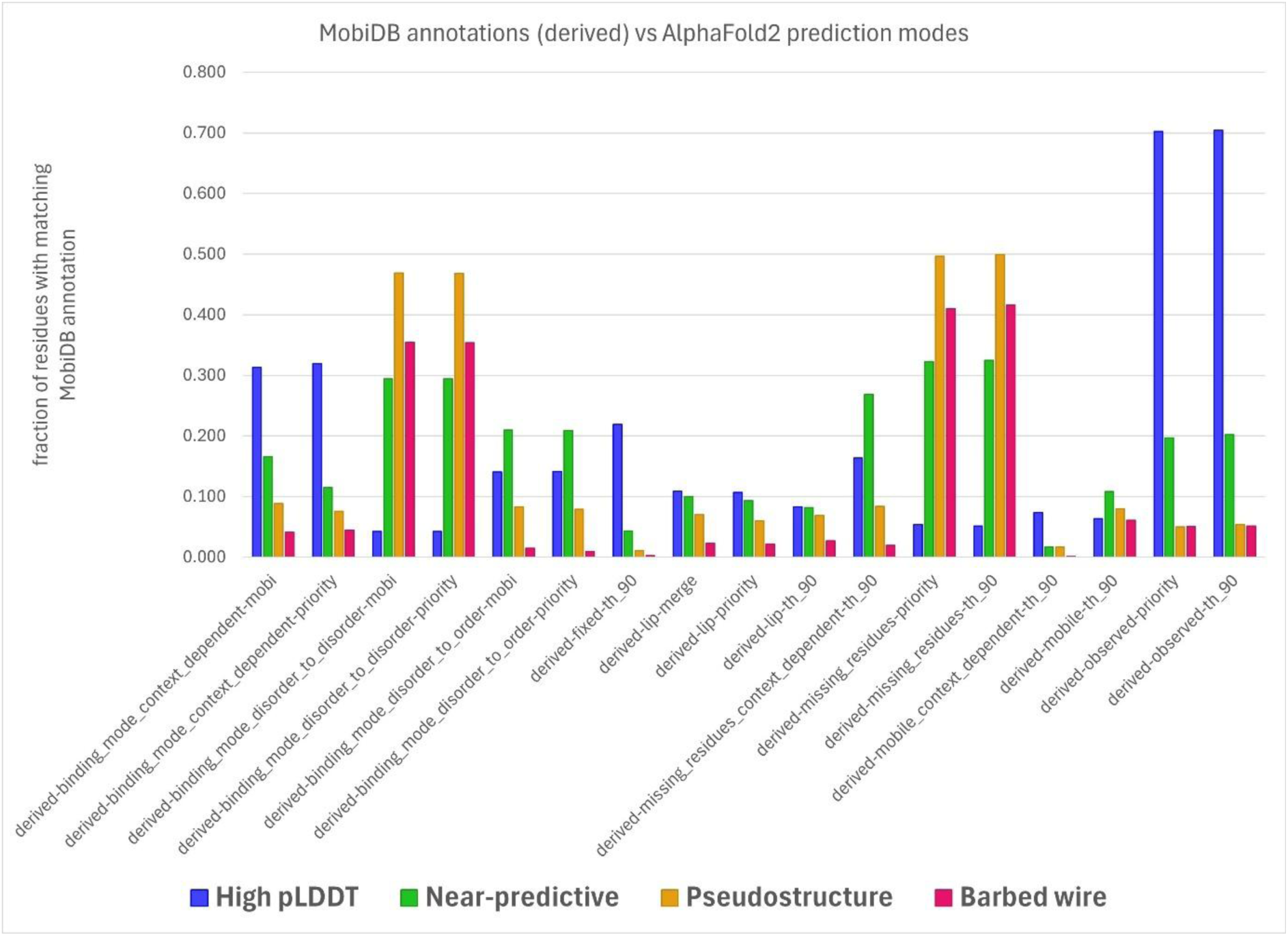
Derived annotations from MobiDB and their relationships with AlphaFold2 prediction modes. Bar height is the fraction of residues from that prediction mode that were marked with the matching MobiDB annotation. Not all sequences are treated with all annotations, and only residues from duly annotated sequences were considered for each annotation. Derived-binding_mode_disorder_to_disorder-mobi and derived-binding_mode_disorder_to_order-mobi are included in Figure 7. Derived-missing_residues-priority is related to residues omitted from experimentally-solved structures. That *near-predictive residues* are frequently missing from their solved structures confirms our difficulty in finding experimentally-solved versions of *near-predictive* regions to check prediction accuracy.

**Figure S4.**
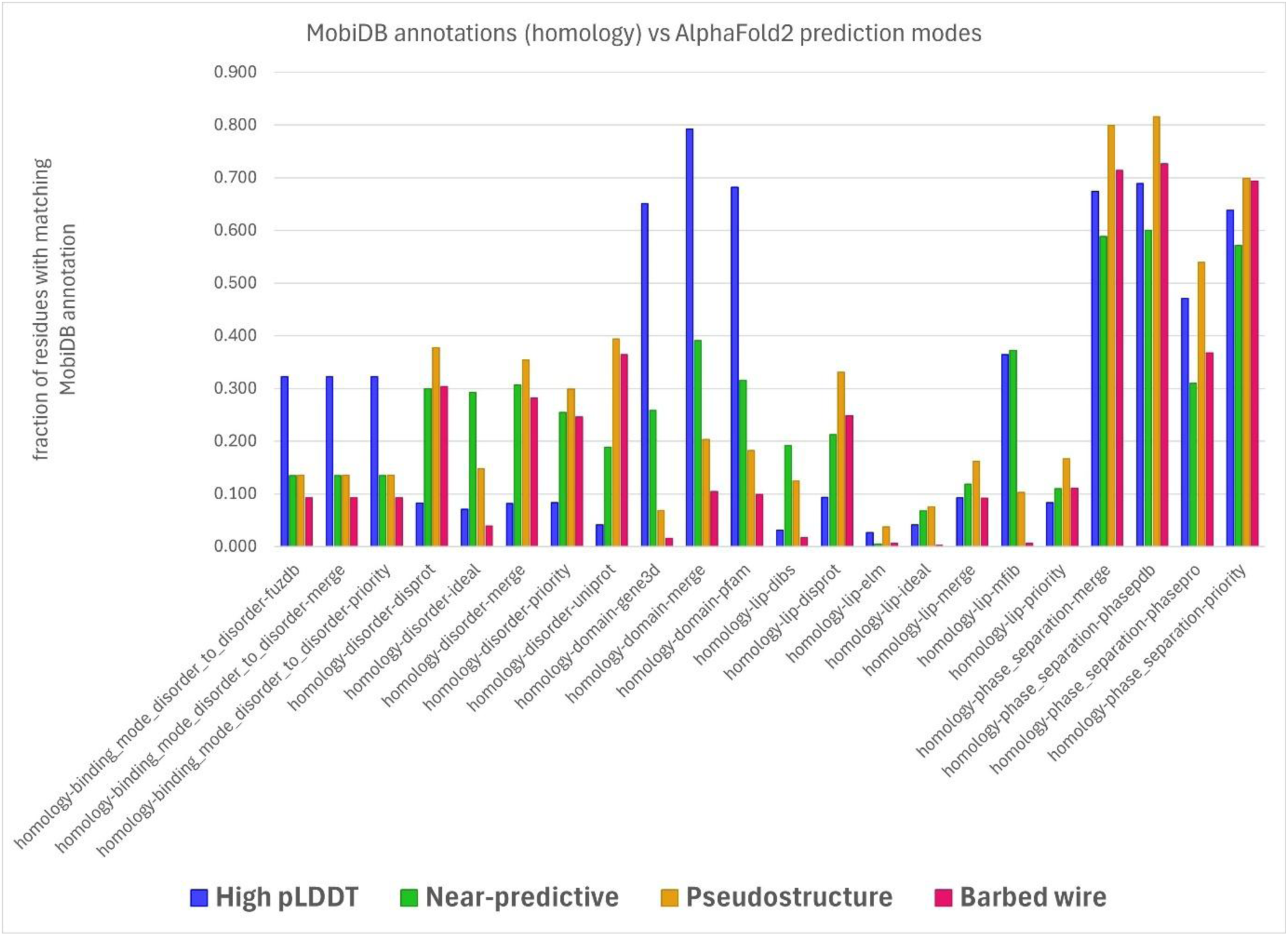
Homology annotations from MobiDB and their relationships with AlphaFold2 prediction modes. Bar height is the fraction of residues from that prediction mode that were marked with the matching MobiDB annotation. Not all sequences are treated with all annotations, and only residues from duly annotated sequences were considered for each annotation. IDEAL is an annotation associated with conditional order; homology-disorder-ideal shows a strong association with *near-predictive*, a lesser association with *pseudostructure*, and very little association with *barbed wire*. This pattern is similar to derived-binding_mode_disorder_to_order-mobi in Figure S3 and supports our conjecture that the *near-predictive* mode is how AlphaFold2 predicts many conditionally ordered IDRs.

**Figure S5.**
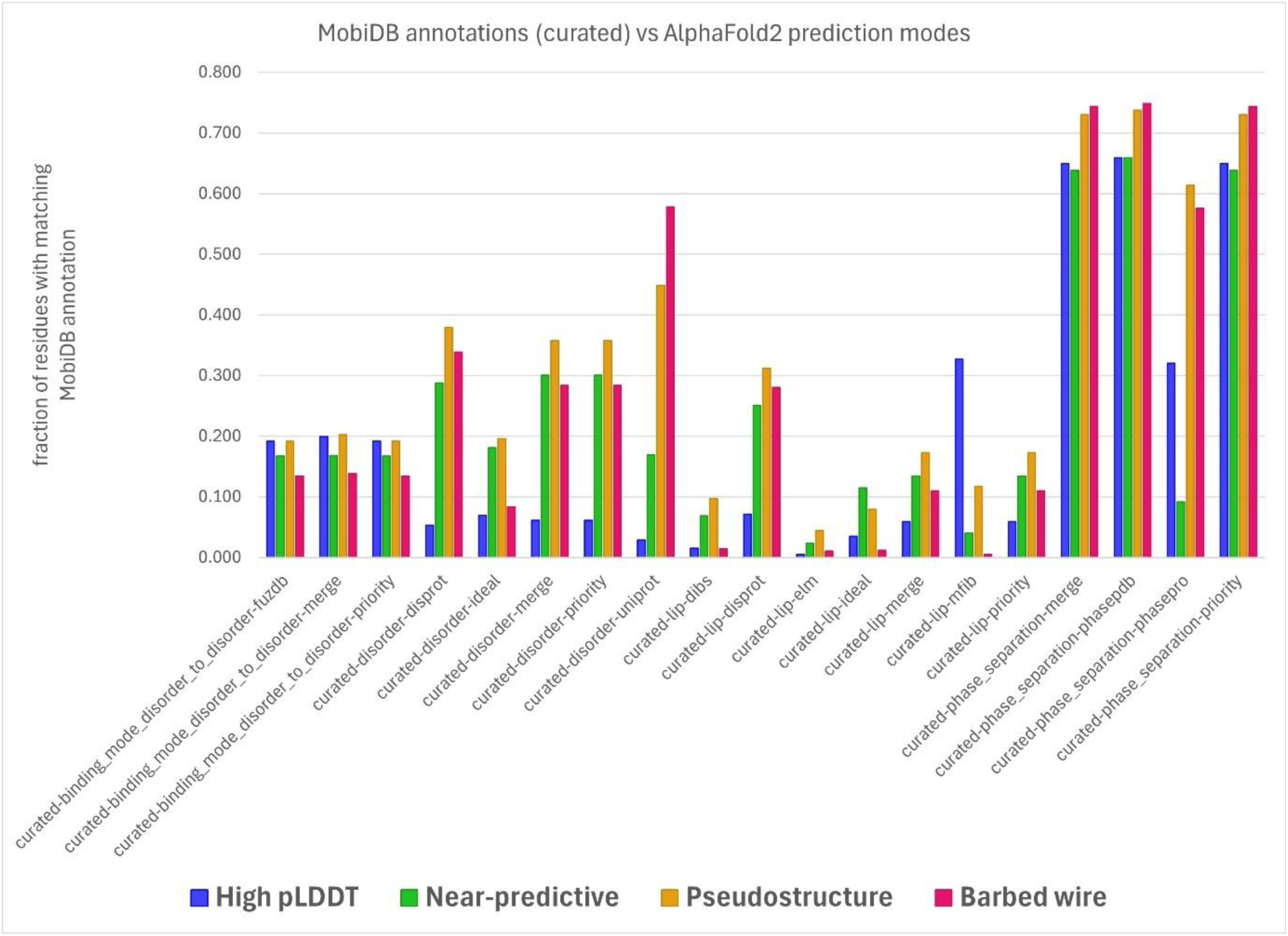
Curated annotations from MobiDB and their relationships with AlphaFold2 prediction modes. Bar height is the fraction of residues from that prediction mode that were marked with the matching MobiDB annotation. Not all sequences are treated with all annotations, and only residues from duly annotated sequences were considered for each annotation. IDEAL is an annotation associated with conditional order. Curated-disorder-ideal shows a more balanced association of *near-predictive* and *pseudostructure* with conditional order than homology-disorder-ideal in Figure S4 above. This may indicate additional complexity in how conditionally folded IDRs manifest in AlphaFold2 predictions, or it may reflect difficulties in IDEAL’s literature-based disorder annotation similar to our own challenges in finding experimentally solved *near-predictive* regions.

## Notes

### Competing Interest Statement

The authors have declared no competing interest.

## References

Abramson, J., Adler, J., Dunger, J., Evans, R., Green, T., Pritzel, A., Ronneberger, O., Willmore, L., Ballard, A. J., Bambrick, J., Bodenstein, S. W., Evans, D. A., Hung, C. C., O’Neill, M., Reiman, D., Tunyasuvunakool, K., Wu, Z., Zemgulyte, A., Arvaniti, E., Beattie, C., Bertolli, O., Bridgland, A., Cherepanov, A., Congreve, M., Cowen-Rivers, A. I., Cowie, A., Figurnov, M., Fuchs, F. B., Gladman, H., Jain, R., Khan, Y. A., Low, C. M. R., Perlin, K., Potapenko, A., Savy, P., Singh, S., Stecula, A., Thillaisundaram, A., Tong, C., Yakneen, S., Zhong, E. D., Zielinski, M., Zidek, A., Bapst, V., Kohli, P., Jaderberg, M., Hassabis, D. & Jumper, J. M. (2024). Nature 630, 493–500.

Adzhubei, A. A., Sternberg, M. J. & Makarov, A. A. (2013). J Mol Biol 425, 2100–2132.

Alderson, T. R., Pritisanac, I., Kolaric, D., Moses, A. M. & Forman-Kay, J. D. (2023). Proc Natl Acad Sci U S A 120, e2304302120.

Baek, M., DiMaio, F., Anishchenko, I., Dauparas, J., Ovchinnikov, S., Lee, G. R., Wang, J., Cong, Q., Kinch, L. N., Schaeffer, R. D., Millan, C., Park, H., Adams, C., Glassman, C. R., DeGiovanni, A., Pereira, J. H., Rodrigues, A. V., van Dijk, A. A., Ebrecht, A. C., Opperman, D. J., Sagmeister, T., Buhlheller, C., Pavkov-Keller, T., Rathinaswamy, M. K., Dalwadi, U., Yip, C. K., Burke, J. E., Garcia, K. C., Grishin, N. V., Adams, P. D., Read, R. J. & Baker, D. (2021). Science 373, 871–876.

Chen, V. B., Davis, I. W. & Richardson, D. C. (2009). Protein Sci 18, 2403–2409.

Conte, A. D., Mehdiabadi, M., Bouhraoua, A., Miguel Monzon, A., Tosatto, S. C. E. & Piovesan, D. (2023). Proteins 91, 1925–1934.

Davis, I. W., Leaver-Fay, A., Chen, V. B., Block, J. N., Kapral, G. J., Wang, X., Murray, L. W., Arendall, W. B., 3rd, Snoeyink, J., Richardson, J. S. & Richardson, D. C. (2007). Nucleic Acids Res 35, W375–383.

Engelbart, D. C. (2023). Augmented Education in the Global Age, pp. 13–29: Routledge.

Fukuchi, S., Sakamoto, S., Nobe, Y., Murakami, S. D., Amemiya, T., Hosoda, K., Koike, R., Hiroaki, H. & Ota, M. (2012). Nucleic Acids Res 40, D507–511.

Jones, D. T. & Thornton, J. M. (2022). Nat Methods 19, 15–20.

Jumper, J., Evans, R., Pritzel, A., Green, T., Figurnov, M., Ronneberger, O., Tunyasuvunakool, K., Bates, R., Zidek, A., Potapenko, A., Bridgland, A., Meyer, C., Kohl, S. A. A., Ballard, A. J., Cowie, A., Romera-Paredes, B., Nikolov, S., Jain, R., Adler, J., Back, T., Petersen, S., Reiman, D., Clancy, E., Zielinski, M., Steinegger, M., Pacholska, M., Berghammer, T., Bodenstein, S., Silver, D., Vinyals, O., Senior, A. W., Kavukcuoglu, K., Kohli, P. & Hassabis, D. (2021). Nature 596, 583–589.

Liebschner, D., Afonine, P. V., Baker, M. L., Bunkoczi, G., Chen, V. B., Croll, T. I., Hintze, B., Hung, L. W., Jain, S., McCoy, A. J., Moriarty, N. W., Oeffner, R. D., Poon, B. K., Prisant, M. G., Read, R. J., Richardson, J. S., Richardson, D. C., Sammito, M. D., Sobolev, O. V., Stockwell, D. H., Terwilliger, T. C., Urzhumtsev, A. G., Videau, L. L., Williams, C. J. & Adams, P. D. (2019). Acta Crystallogr D Struct Biol 75, 861–877.

Lin, Z., Akin, H., Rao, R., Hie, B., Zhu, Z., Lu, W., Smetanin, N., Verkuil, R., Kabeli, O., Shmueli, Y., Dos Santos Costa, A., Fazel-Zarandi, M., Sercu, T., Candido, S. & Rives, A. (2023). Science 379, 1123–1130.

Lovell, S. C., Davis, I. W., Arendall, W. B., 3rd, de Bakker, P. I., Word, J. M., Prisant, M. G., Richardson, J. S. & Richardson, D. C. (2003). Proteins 50, 437–450.

Matthews, B. (1972). Macromolecules 5, 818–819.

Moriarty, N. W., Tronrud, D. E., Adams, P. D. & Karplus, P. A. (2014). The FEBS journal 281, 4061–4071.

Necci, M., Piovesan, D., Predictors, C., DisProt, C. & Tosatto, S. C. E. (2021). Nat Methods 18, 472–481.

Piovesan, D., Del Conte, A., Mehdiabadi, M., Aspromonte, M. C., Blum, M., Tesei, G., von Bulow, S., Lindorff-Larsen, K. & Tosatto, S. C. E. (2025). Nucleic Acids Res 53, D495–D503.

Piovesan, D., Monzon, A. M. & Tosatto, S. C. (2022). Protein Science 31, e4466.

Prisant, M. G., Williams, C. J., Chen, V. B., Richardson, J. S. & Richardson, D. C. (2020). Protein Sci 29, 315–329.

Ramachandran, G. N., Ramakrishnan, C. & Sasisekharan, V. (1963). J Mol Biol 7, 95–99.

Richardson, J. S. (2000). Nat Struct Biol 7, 624–625.

Richardson, J. S., Williams, C. J., Chen, V. B., Prisant, M. G. & Richardson, D. C. (2023). Acta Crystallogr D Struct Biol 79, 1071–1078.

Sasisekharan, V. (1959). Acta Crystallographica 12, 897–903.

Sehnal, D., Bittrich, S., Deshpande, M., Svobodova, R., Berka, K., Bazgier, V., Velankar, S., Burley, S. K., Koca, J. & Rose, A. S. (2021). Nucleic Acids Res 49, W431–W437.

Tam, C. & Iwasaki, W. (2023). Proteomics 23, e2300176.

Terwilliger, T. C., Afonine, P. V., Liebschner, D., Croll, T. I., McCoy, A. J., Oeffner, R. D., Williams, C. J., Poon, B. K., Richardson, J. S., Read, R. J. & Adams, P. D. (2023). Acta Crystallogr D Struct Biol 79, 234–244.

Terwilliger, T. C., Liebschner, D., Croll, T. I., Williams, C. J., McCoy, A. J., Poon, B. K., Afonine, P. V., Oeffner, R. D., Richardson, J. S., Read, R. J. & Adams, P. D. (2024). Nat Methods 21, 110–116.

Thoms, M., Buschauer, R., Ameismeier, M., Koepke, L., Denk, T., Hirschenberger, M., Kratzat, H., Hayn, M., Mackens-Kiani, T., Cheng, J., Straub, J. H., Sturzel, C. M., Frohlich, T., Berninghausen, O., Becker, T., Kirchhoff, F., Sparrer, K. M. J. & Beckmann, R. (2020). Science 369, 1249–1255.

Thornton, J. M., Laskowski, R. A. & Borkakoti, N. (2021). Nature Medicine 27, 1666–1669.

Tunyasuvunakool, K., Adler, J., Wu, Z., Green, T., Zielinski, M., Zidek, A., Bridgland, A., Cowie, A., Meyer, C., Laydon, A., Velankar, S., Kleywegt, G. J., Bateman, A., Evans, R., Pritzel, A., Figurnov, M., Ronneberger, O., Bates, R., Kohl, S. A. A., Potapenko, A., Ballard, A. J., Romera-Paredes, B., Nikolov, S., Jain, R., Clancy, E., Reiman, D., Petersen, S., Senior, A. W., Kavukcuoglu, K., Birney, E., Kohli, P., Jumper, J. & Hassabis, D. (2021). Nature 596, 590–596.

Varadi, M., Bertoni, D., Magana, P., Paramval, U., Pidruchna, I., Radhakrishnan, M., Tsenkov, M., Nair, S., Mirdita, M., Yeo, J., Kovalevskiy, O., Tunyasuvunakool, K., Laydon, A., Zidek, A., Tomlinson, H., Hariharan, D., Abrahamson, J., Green, T., Jumper, J., Birney, E., Steinegger, M., Hassabis, D. & Velankar, S. (2024). Nucleic Acids Res 52, D368–D375.

Wang, W., Gong, Z. & Hendrickson, W. A. (2025). Acta Crystallogr D Struct Biol 81, 4–21.

Williams, C. J., Headd, J. J., Moriarty, N. W., Prisant, M. G., Videau, L. L., Deis, L. N., Verma, V., Keedy, D. A., Hintze, B. J., Chen, V. B., Jain, S., Lewis, S. M., Arendall, W. B., 3rd, Snoeyink, J., Adams, P. D., Lovell, S. C., Richardson, J. S. & Richardson, D. C. (2018). Protein Sci 27, 293–315.

Wilson, C. J., Choy, W. Y. & Karttunen, M. (2022). Int J Mol Sci 23.

